# Proteomic analysis reveals dysregulation of peripheral blood neutrophils in patients with Multiple Sclerosis

**DOI:** 10.1101/2024.05.29.596498

**Authors:** Katie J Smith, Zachary Lim, Sonja Vermeren, Veronique E. Miron, Sarah Dimeloe, Donald J Davidson, Anna Williams, Emily Gwyer Findlay

## Abstract

Multiple Sclerosis (MS) is a complex auto-inflammatory disease affecting the brain and spinal cord, which results in axonal de-myelination and symptoms including fatigue, pain, and difficulties with vision and mobility. The involvement of the immune system in the pathology of MS is well established, particularly the adaptive T cell response, and there has been a particular focus on the IL-17-producing subset of Th17 cells and their role in driving disease. However, the importance of innate immune cells has not been so well characterised. Here we focused on neutrophils, which are innate immune cells and rapid responders to inflammation, and which have recently been linked to other chronic autoimmune conditions. Multiple strands of evidence in patients with MS and in mice with the experimental autoimmune encephalomyelitis MS model suggest neutrophils may play a role in driving MS inflammation. Here, we performed proteomic analysis on neutrophils from patients with MS and healthy donors, revealing striking differences. In particular, granule proteins were significantly more abundant in the MS neutrophils compared to the healthy controls, with a particular over-abundance of proteins in primary and secondary granules. In addition, members of the MAVS signalling pathway were differently regulated compared to healthy donor cells. Finally, we find that MS neutrophils do not suppress T cell activation equivalently to healthy neutrophils, and in particular are unable to suppress expression of CD161 on the T cells, indicative of Th17 differentiation. We propose that neutrophil dysregulation in MS may contribute to dysfunctional T cell responses.

## Introduction

Multiple Sclerosis (MS) is an auto-inflammatory disease affecting the central nervous system. The development and pathology of MS are complex, with early inflammation leading to later neuro-degeneration, axonal demyelination and axonal loss. The contributions of adaptive immunity such as T cells (in particular Th1 and Th17 cells and their cytokines) to the inflammatory stage of MS development are well-established ^1–7^, but other contributors to inflammation in early MS are less understood. In particular, the role of innate immune cells such as neutrophils is unclear.

Neutrophils are the most abundant leukocyte in human blood and the first incoming responders to infectious and sterile inflammation. Owing to their rapid recruitment during the initiation of inflammation, short lifespan, and rapid clearance following death, the role of neutrophils in driving chronic diseases such as MS was long underestimated. More recently, however, neutrophil half-life estimations have been considerably extended ^8^; continual neutrophilia has been found to be a feature of other chronic autoimmune and autoinflammatory conditions ^9,10^; and their interactions with adaptive immune cells such as T cells has been revealed to be specific and complex ^11–16^. An interest in the role of neutrophils in long-term autoimmune diseases like MS is therefore developing.

It is now well established that an increased systemic neutrophil to lymphocyte ratio is a marker of MS disease activity ^17–20^. Locally, neutrophils have been found in active brain lesions ^12,21^. Not only the frequency of neutrophils but also their phenotype is changed with disease - peripheral blood neutrophils from patients with MS are typically more activated, produce more reactive oxygen species and produce neutrophil extracellular traps (NETs) more rapidly than those from healthy donors ^17,22^.

Correlative studies suggest reducing neutrophil numbers would be beneficial in disease. A positive response to treatment correlates with neutropenia ^23^, and in contrast, GCSF therapy (which stimulates production and emigration of immature neutrophils from the bone marrow) worsens disease exacerbations ^24^.

The mouse model of the inflammatory stage of MS, experimental autoimmune encephalomyelitis (EAE), also shows strong pathogenic roles for these cells. Mice depleted of neutrophils do not develop disease ^25,26^.

Neutrophils isolated from the spinal cord during disease promote the activation and maturation of antigen presenting cells ^25^. We have also previously shown that neutrophils in the spinal cord and brain of affected mice direct a pathogenic T cell response through release of the granular antimicrobial peptide cathelicidin ^12^.

All of this work demonstrates a clear need for further investigation into neutrophils in patients with MS – yet no unbiased screening of these cells has yet been performed. Here, we undertook such an approach through proteomic analysis of rapidly-isolated neutrophils from patients with MS and healthy donors. Our results show a profound and consistent dysregulation of peripheral blood neutrophils in patients diagnosed with MS, with neutrophils appearing more mature, having consistently more abundant antimicrobial peptides, and altered expression of members of the mitochondrial antiviral-signalling protein (MAVS) signalling pathway. Co-culture of these neutrophils with T cells showed that T cells contacting healthy neutrophils are suppressed in their activation and less likely to express markers of Th17 differentiation, while T cells contacting MS neutrophils do not show this suppression. This suggests a route by which dysregulated neutrophils may lead to differences in adaptive immune responses in autoimmune disease.

## Methods

### Patients and ethical agreements

The study was approved by Lothian Bioresource under agreement SR1323. Patients with MS attending the Anne Rowling Regenerative Neurology Clinic (Edinburgh, Scotland) for post-diagnosis clinical meetings were enrolled in the study, with full written consent, by clinical staff. Exclusion criteria were: under the age of 18; over the age of 65; and on any disease-modifying MS immune-modulatory drug therapies. Healthy donor blood was collected under ethical agreement 21-EMREC-041, with full written consent (Table 1).

**Table 1:** Proteomics donor information table. MS, multiple sclerosis; HD, healthy donor; RRMS, relapsing-remitting multiple sclerosis; SPMS, secondary progressive multiple sclerosis; PPMS, primary progressive multiple sclerosis

| Donor | Age | Sex | Diagnosis |
| --- | --- | --- | --- |
| MS1 | 31-40 | F | RRMS |
| MS2 | 61-70 | F | RRMS |
| MS3 | 31-40 | F | SPMS |
| MS4 | 31-40 | F | RRMS |
| MS5 | 41-50 | F | RRMS |
| MS6 | 51-60 | M | RRMS |
| MS7 | 51-60 | F | PPMS |
| MS8 | 51-60 | F | SPMS |
| HD1 | 41-50 | F | - |
| HD2 | 21-30 | F | - |
| HD3 | 41-50 | F | - |
| HD4 | 41-50 | M | - |
| HD5 | 21-30 | F | - |
| HD6 | 41-50 | F | - |
| HD7 | 31-40 | F | - |
| HD8 | 21-30 | F | - |

### Isolation of neutrophils and T cells

Highly purified neutrophils and CD3^+^ T cells were immediately isolated from fresh blood using immunomagnetic negative separation. The Stem Cell Technologies direct neutrophil isolation kit (StemCell Technologies, #17957) and Stem cell Human T cell isolation kit (StemCell Technologies, #17951) were used respectively, as per manufacturer’s instructions.

### Flow cytometry

Cells were stained with either 1 μg/ml DAPI (Invitrogen #D1306) or 1:1000 live/dead yellow (Invitrogen #L-34959) diluted in 1XPBS, or 1:150 Zombie NIR Fixable Viability Kit (Biolegend, #423105) diluted in 1XPBS for 20 minutes at room temperature, protected from light. Surface markers (listed below) were stained for 30 minutes at 4°C in 1XPBS, protected from light. Samples were run on a BD Biosciences LSR Fortessa cytometer and data analysed using FlowJo software, version 10 (BD Biosciences).

Antibodies used include: CD3 FITC (clone HIT3a, Biolegend, #300305, 1:200), CD4 PeCy7 (clone A161A1, Biolegend, #357409, 1:200), CD8 APC (clone SK1, Biolegend, #344721, 1:200), CD62L PeDazzle (clone DREG-56, Biolegend, #304805, 1:200), CD62L AF488 (clone DREG-56, Biolegend, #304816, 1:150), CD161 APC (clone HP-3G10, Biolegend, #339911, 1:200), CD11b APC/fire (clone CBRM1/5, Biolegend, #301419, 1:150), CD44 BV510 (clone IM7, Biolegend, #103049, 1:400).

### Preparation of neutrophils for proteomic analysis

Sample preparations and mass spectrometry was performed as a service by Fingerprints Proteomics facility, University of Dundee.

Following isolation, neutrophils were centrifuged at 10,000g for 8 minutes, supernatant removed and discarded, and the neutrophil pellet was snap-frozen and stored at -80°C until mass spectrometry was carried out. Pellets were then lysed in buffer containing 5% SDS and 50 mM triethylammonium bicarbonate and sonicated for 30 seconds on/ 30 seconds off at 50% amplitude. 20mM dithiothreitol was added and cells were boiled at 95 °C for 10 minutes. Once cooled, 40mM iodoacetamide was added for 30 minutes at room temperature. Proteins were isolated with S-TRAP columns. Cells were digested with trypsin overnight at 37°C before more trypsin was added for 6 hours and proteins collected by elution from the columns with 50mM triethylammonium bicarbonate, 0.2% aqueous formic acid and 50% aqueous acetonitrile with 0.2% formic acid. Protein concentration was determined using the EZQ protein quantitation kit (Invitrogen, #R33200).

### Liquid chromatography and mass spectrometry (LC/MS)

Neutrophil proteins were quantified and normalised using a microBSA assay. 1.5 µg of peptide was analysed per sample. Samples were injected onto a nanoscale C18 reverse-phase chromatography system (UltiMate 3000 RSLC nano, Thermo Scientific) then electrosprayed into an Q Exactive HF-X Mass Spectrometer (Thermo Scientific). For liquid chromatography buffers were as follows: buffer A (0.1% formic acid in Milli-Q water (v/v)) and buffer B (80% acetonitrile and 0.1% formic acid in Milli-Q water (v/v). Sample were loaded at 10 μL/min onto a trap column (100 μm × 2 cm, PepMap nanoViper C18 column, 5 μm, 100 Å, Thermo Scientific) equilibrated in 0.1% trifluoroacetic acid (TFA). The trap column was washed for 3 minutes at the same flow rate with 0.1% TFA then switched in-line with a Thermo Scientific, resolving C18 column (75 μm × 50 cm, PepMap RSLC C18 column, 2 μm, 100 Å). The peptides were eluted from the column at a constant flow rate of 300 nl/min with a linear gradient from 3% buffer B to 6% buffer B in 5 min, then from 6% buffer B to 35% buffer B in 115 minutes, and finally to 80% buffer B within 7 minutes. The column was then washed with 80% buffer B for 4 minutes and re-equilibrated in 35% buffer B for 5 minutes. Two blanks were run between each sample to reduce carry-over. The column was kept at a constant temperature of 40 °C.

The data was acquired using an easy spray source operated in positive mode with spray voltage at 1.950 kV, and the ion transfer tube temperature at 250 °C. The MS was operated in DIA mode. A scan cycle comprised a full MS scan (m/z range from 350-1650), with RF lens at 40%, AGC target set to custom, normalised AGC target at 300, maximum injection time mode set to custom, maximum injection time at 20 ms and source fragmentation disabled. MS survey scan was followed by MS/MS DIA scan events using the following parameters: multiplex ions set to false, collision energy mode set to stepped, collision energy type set to normalized, HCD collision energies set to 25.5, 27 and 30, orbitrap resolution 30000, first mass 200, RF lens 40, AGC target set to custom, normalized AGC target 3000, maximum injection time 55 ms.

Data analysis was carried out using Spectonaut (version 16.2.220903.53000, Biognosys, AG). The directDIA workflow, using the default settings (BGS Factory Settings) with the following modifications was used: decoy generation set to mutated; Protein LFQ Method was set to QUANT 2.0 (SN Standard) and Precursor Filtering set to Identified (Qvalue); Precursor Qvalue Cutoff and Protein Qvalue Cutoff (Experimental) set to 0.01; Precursor PEP Cutoff set to 0.1 and Protein Qvalue Cutoff (Run) set to 0.05. The data were normalized on the global median. For the Pulsar search the settings were: maximum of 2 missed trypsin cleavages; PSM, Protein and Peptide FDR levels set to 0.01; scanning range set to 300-1800 m/z and Relative Intensity (Minimum) set to 5%; cysteine carbamidomethylation set as fixed modification and acetyl (Protein N-term), deoxidation (methionine, tryptophan), deamidation (asparagine, glutamine) and oxidation of methionine set as variable modifications. The database used was H.sapiens proteome downloaded from 6niport.org on 2021-05-11 (77027 entries).

### Data

The proteomic dataset collected is available at FigShare, https://doi.org/10.6084/m9.figshare.24948342.v2

### *In vitro* co-culture system

75,000 T cells and 225,000 neutrophils were cultured together in one well of a 96-well plate. MS patient and healthy donor T cells were plated alone, or co-cultured with neutrophils, with 0.5µl/ml CD3/CD28/CD2 T cell activator (STEMCELL Technologies, #10970). Cells were incubated at 37°C for 24h.

### R analysis

The volcano plot (Fig2A) was generated in R using the ggplot2 (v3.5.0) and ggrepel (v0.9.5) package with default parameters. Upregulated genes were defined as mlog10p value of > 1.3 and a log 2-fold change value of > 1 and downregulated genes were defined as mlog10p value of > 1.3 and a log 2-fold change value of < -1. Pathway analysis (Fig2D) was performed in R using the clusterProfiler (v4.10.1) package with the enrichGO function with a p-value cut off as 0.05.

## Results

No unbiased analysis of neutrophils in MS has yet been performed, although there is mounting evidence that the cells are dysregulated in disease. To perform these experiments, we collected peripheral blood from patients with MS and from control donors.

### Peripheral blood neutrophils are activated in patients with MS

There were no differences in total neutrophil number in peripheral blood between the groups (Fig1A). First, we analysed the cell surface phenotype of the neutrophils using flow cytometry for indicators of increased activation. Neutrophils from patients with MS showed lower expression of surface CD62L (Fig1B, C) and higher CD11b expression (Fig1D, E) than healthy donor cells, both indicative of increased activation. These data suggested that circulating neutrophils are activated in patients with MS, supporting a previous report^17^, which also demonstrated the activation of MS neutrophils.

**Figure 1:**
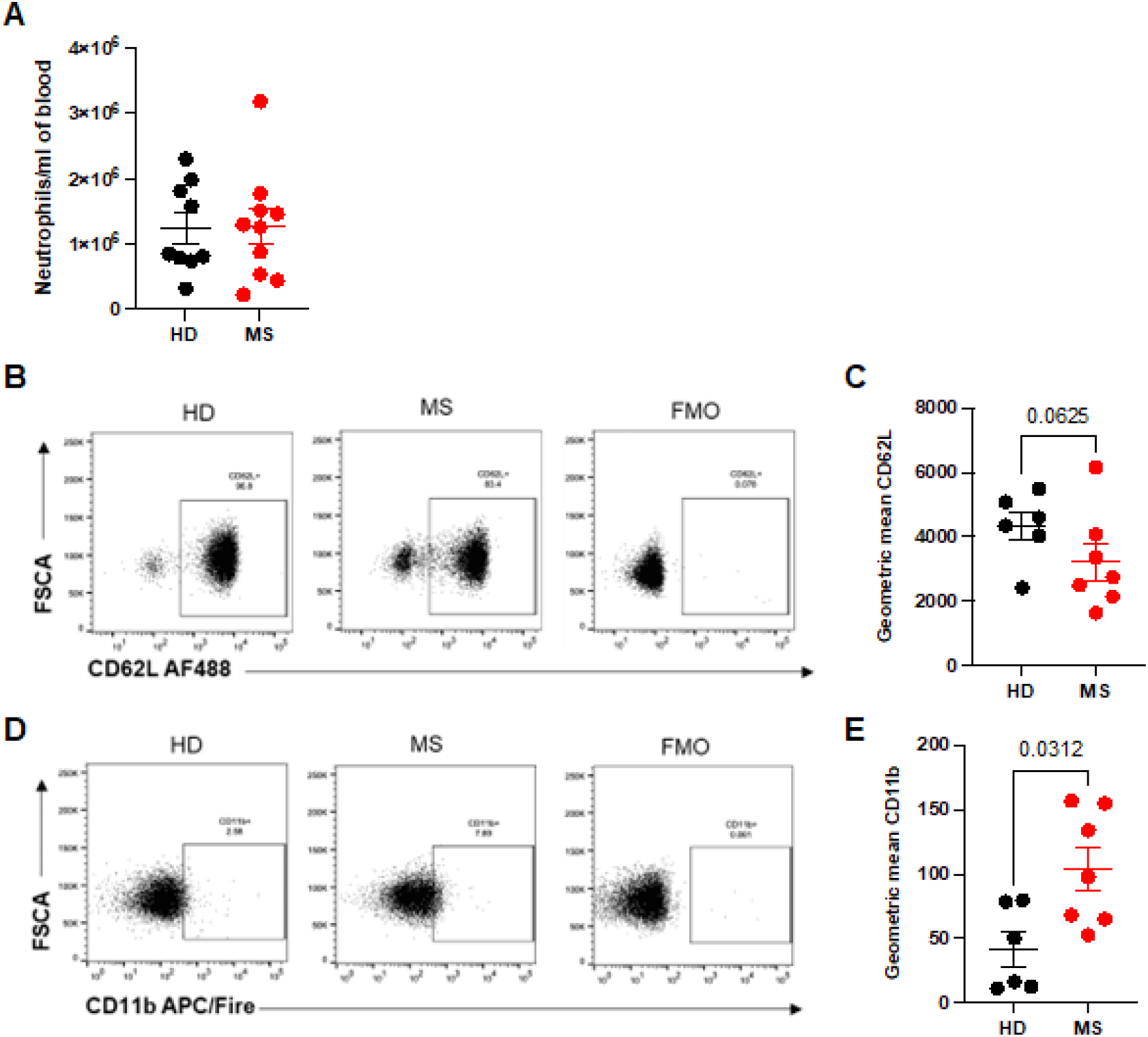
Peripheral blood neutrophils from patients with multiple sclerosis are more activated than cells from healthy donors. Peripheral blood was drawn from patients with multiple sclerosis (MS, red) and healthy donors (HD, black). Neutrophils were isolated immediately via negative magnetic selection and (A) cell counts performed using a haemocytometer. (B-E) Neutrophil activation was measured by CD62L loss and CD11b gain. HD – healthy donor; MS – multiple sclerosis; FMO – fluorescent minus one. N values: A-E – 6–10 donors. Statistical test: C,E – Wilcoxon matched-pairs test.

These initial results confirmed the importance of performing an unbiased, larger-scale analysis of these cells. Since neutrophils have a comparatively low transcriptional activity, with many of their most abundant proteins stored in granules, we opted to use proteomics rather than RNA sequencing for their analysis; indeed, a number of recent studies have demonstrated the value of this technique in other diseases ^27–29^. We therefore progressed by performing mass spectrometry analysis of proteins on frozen neutrophil pellets from our 16 samples.

### Neutrophils from patients with MS are activated and mature

A total of 3890 unique proteins were identified in the samples, with differences clearly observable between the two groups (Fig2A). As our initial studies had demonstrated that MS neutrophils were characterised by an activated phenotype, we first analysed the differential expression of markers of neutrophil inflammation and maturation. Of particular interest, the surface protein CD10 (membrane metalloendopeptidase) was up-regulated in MS neutrophils (146% of HD value on average, Fig2B). In addition, the proliferation marker PCNA (proliferating cell nuclear antigen) was down-regulated in these samples (64% of average HD value, Fig2C). This expression pattern suggests increased priming and maturation of neutrophils in the periphery of patients with MS. Interestingly, the two patients with secondary progressive MS (blue symbols) did not differentiate from the patients with relapsing remitting MS (red symbols) or primary progressive MS (grey symbol). Pathway analysis of differentially expressed proteins showed significant alterations in anti-viral responses and response to oxidative stress, as well as demonstrating altered intensity of proteins related to translation and cell-cell adhesion (Fig2D).

**Figure 2:**
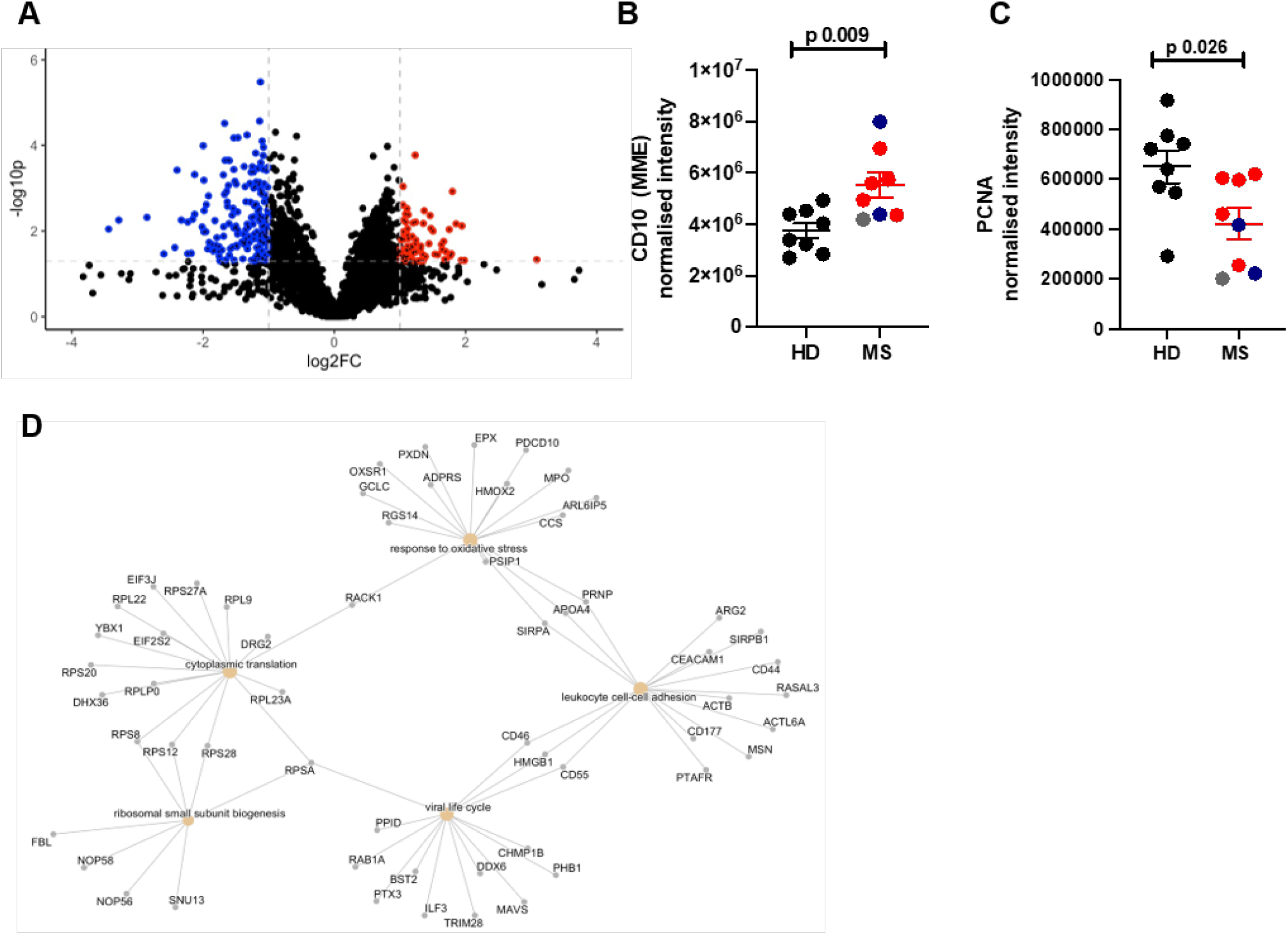
Neutrophils from patients with MS are more mature. Peripheral blood was drawn from patients (MS) and healthy donors (HD). Neutrophils were isolated immediately via negative magnetic selection, centrifuged and cell pellets were snap-frozen. Cell pellets were analysed via LC/MS for protein abundance. (A) Unbiased volcano analysis of upregulated (red) and downregulated (blue) proteins in MS neutrophils compared to healthy donor cells defined by - log 10pvalue of <1.3 and a Iog2fold change of > 1 or < -1, represented by the dotted lines. (B, C) Neutrophil maturity was assessed via CD10 and PCNA abundance in RRMS (red), SPMS (blue), PPMS (grey) and HD (black). (D) Pathway analysis of differentially expressed proteins in MS neutrophils compared to healthy donor neutrophils. HD – healthy donor; MS – multiple sclerosis; FC – fold change. N values: A-D –8. Statistical test: B, C – unpaired t test with Welch’s correction.

### Antimicrobial peptides and proteins are more abundant in MS neutrophils

Patients with MS are at increased risk of infection, even if they are not on any disease modifying therapy ^30–32^. We hypothesised that one reason for this may be that their neutrophils have impaired responses to pathogens. We therefore analysed the dataset for proteins relating to innate immune and anti-infection processes.

We noted first that antimicrobial peptides and proteins in MS neutrophils were significantly more abundant compared to HD cells. Cathelicidin (CATH), lactotransferrin (LTF), and multiple alpha defensins (DEFA1, DEFA3, DEFA4) were all present at higher levels in MS neutrophils than healthy controls (Fig3A-E).

**Figure 3:**
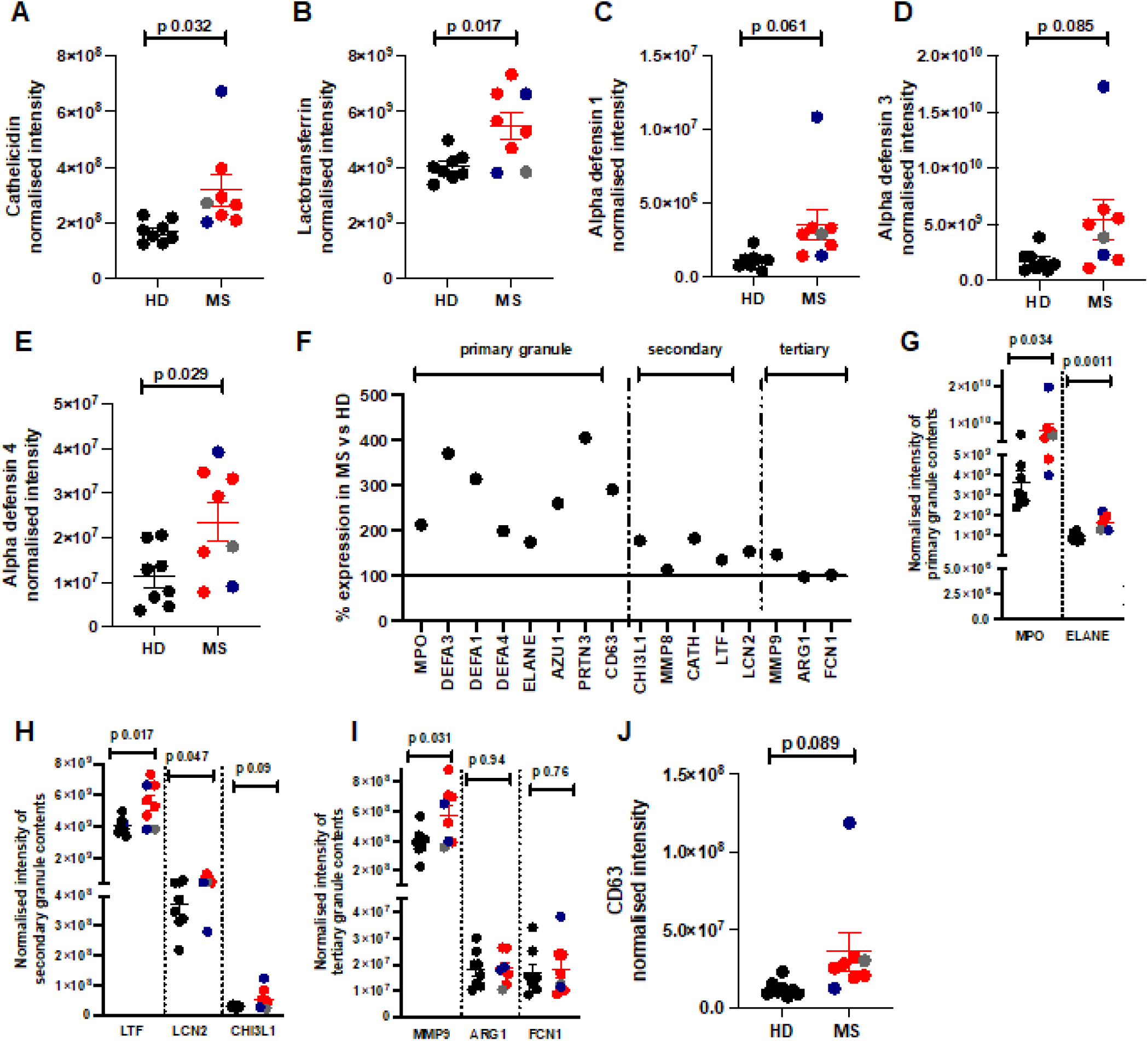
Increased abundance of primary and secondary granule proteins in MS neutrophils. Peripheral blood was drawn from patients (RRMS, red, SPMS, blue or PPMS, grey) and healthy donors (HD, black). Neutrophils were isolated immediately via negative magnetic selection, centrifuged and cell pellets were snap-frozen. (A-E, G-K) Cell pellets were analysed via LC/MS for protein abundance. (F) granule protein abundance in MS neutrophils as a percentage of healthy donor abundance, line at 100% expression. HD – healthy donor; MS – multiple sclerosis. N values: A-K –8. Statistical test: unpaired t test with Welch’s correction.

Many antimicrobial peptides and proteins are stored in the cytoplasmic granules, of which there are three major types – primary, secondary and tertiary. We therefore extended our analysis to look at the granule protein composition in neutrophils from each group. Interestingly, there were large changes (% alteration shown in Fig3F, individual values in Fig3G-I) which differed for different constituent proteins/peptides across the various types of granules. Primary granule contents were significantly more abundant in MS neutrophils compared to healthy neutrophils (Fig3F, G), with an average of all primary granule contents at 280% of the abundance in healthy donor cells, as demonstrated with myeloperoxidase (MPO) and elastase (ELANE) (Fig3G). The secondary granule protein abundance was also increased compared to healthy cells (Fig 3F, H), although less so than the primary granules, with a median abundance of 154% of the healthy value, demonstrated here with lactotransferrin (LTF), lipocalin 2 (LCN2) and chitinase 3-like protein 1 (CHI3L1) (Fig3H). Interestingly, the tertiary granule contents were not more abundant in the MS neutrophils than the healthy cells (Fig3F, I), with the exception of MMP9 – shown here also are arginase 1 (ARG1) and ficolin 1 (FCN1).

An increase in abundance of granule proteins is intriguing as MS neutrophils are more activated, therefore we would have hypothesised that they undergo increased degranulation and consequently contain lower concentrations of granule proteins. In support of this, others have shown^17,22^ that CD63 expression, which can be used as a marker for release of primary granules ^33^, is increased on MS neutrophils compared to healthy controls. We therefore examined expression of CD63 in our dataset. We found it to be elevated in the MS neutrophils, although very variable between patients, with a mean of 292% of the expression seen in healthy neutrophils, (Fig3J), agreeing with this previous work. This data therefore suggests a combination of altered granule composition and degranulation rates in the MS neutrophils.

### Anti-infection responses are increased in MS neutrophils

As our data are suggestive of dysregulated degranulation in MS neutrophils, we next wondered if other anti-microbial responses were altered in MS neutrophils, such as reactive oxygen species. Firstly, ‘response to oxidative stress’ was found to be altered in the pathway analysis (Fig2D). Secondly, we had noted previously that PCNA is expressed at lower levels in MS neutrophils (Fig2C). PCNA promotes the production of reactive oxygen species (ROS) ^34^ and so we hypothesised that MS neutrophils would produce lower concentrations of ROS. To test this hypothesis, we examined the expression of proteins involved in the generation of ROS: p47phox (NCF1), p67phox (NCF2), Gp91phox (NOX2), p22phox (CYBA) and p40phox (NCF4). gp91phox was not found in the dataset. Of the others, p47phox and p67phox showed no difference between patients and healthy donors, while p22phox was significantly increased and p40phox was decreased in MS neutrophils (Fig4A-D).

**Figure 4:**
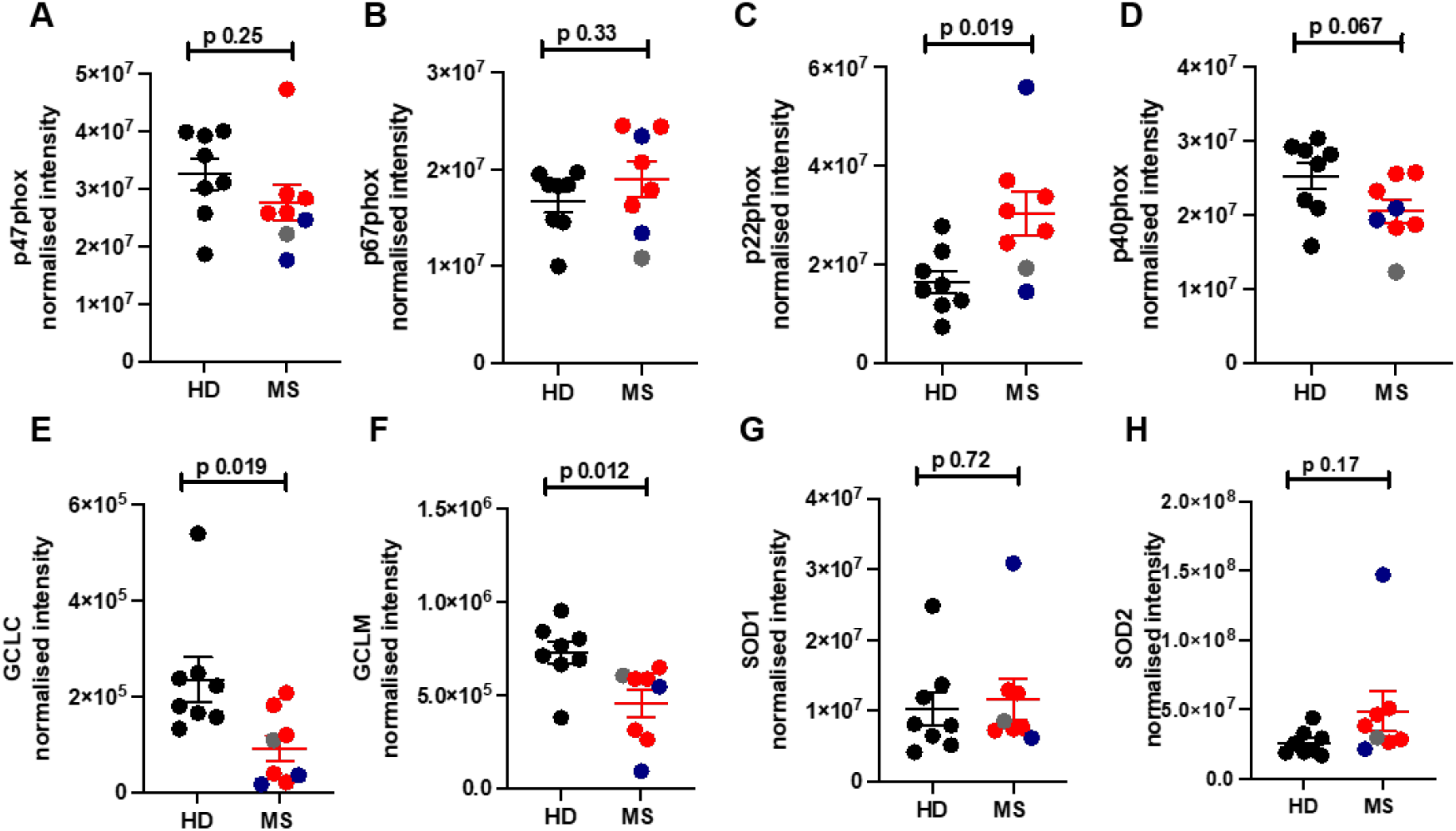
Alterations in reactive oxygen pathways in MS neutrophils. Peripheral blood was drawn from patients (RRMS, red, SPMS, blue or PPMS, grey) and healthy donors (HD, black). Neutrophils were isolated immediately via negative magnetic selection, centrifuged and cell pellets were snap-frozen. (A-l) Cell pellets were analysed via LC/MS for protein abundance. HD – healthy donor; MS – multiple sclerosis. N values: A-l –8. Statistical test: A-l – unpaired t test with Welch’s correction

To further investigate whether reactive oxygen pathways were altered in neutrophils from patients, we analysed the abundance of glutamate-cysteine ligase (GCLC), glutamate-cysteine ligase regulatory subunit (GCLM), glutathione peroxidase 3 (GPX3), superoxide dismutase 1 (SOD1) and superoxide dismutase 2 (SOD2) (Fig4E-H). Interestingly, GCLC ligase (Fig4E) and GCLM (Fig4F) were significantly down-regulated in the MS neutrophils. No other proteins were altered between the two groups.

Next, we had seen in the pathway analysis that MAVS came up as a key protein altered in abundance (Fig2D). We noted in the proteomic dataset that members of this MAVS signalling pathway were particularly affected in the patient cells (Fig5A). This is a key pathway through which interferons are produced, although it is not well characterised in neutrophils. In the MS neutrophils there was a large and significant decrease in the MAVS protein (Fig5B) and increases in the abundance of IFIT3, RIG-I (DDX58), DHX36 and PP6C (Fig 5C-F).

**Figure 5:**
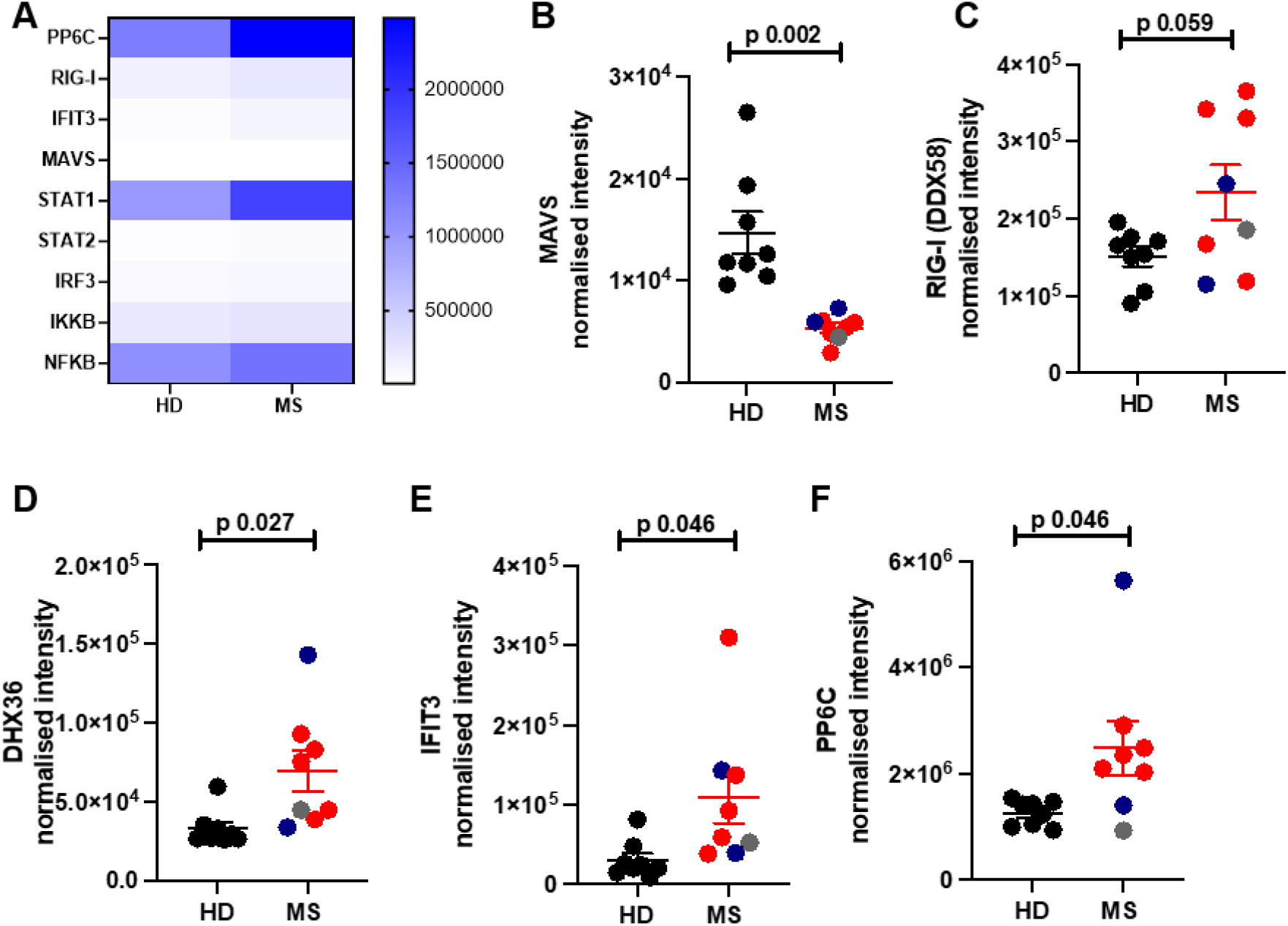
Altered abundance of MAVS pathway members in MS neutrophils. Peripheral blood was drawn from patients (RRMS, red, SPMS, blue or PPMS, grey) and healthy donors (HD, black). Neutrophils were isolated immediately via negative magnetic selection, centrifuged and cell pellets were snap-frozen. (A-F) Cell pellets were analysed via LC/MS for protein abundance. HD – healthy donor; MS – multiple sclerosis. N values: A-F –8. Statistical test: B-F – unpaired t test with Welch’s correction.

### MS neutrophils have altered carbohydrate and lipid metabolism

We were interested in how proteins relating to metabolism and metabolic signalling pathways were altered in MS neutrophils, since others have demonstrated altered neutrophil metabolism in chronic inflammatory conditions such as rheumatoid arthritis ^10^. Neutrophil metabolism has generally been considered to be predominantly glycolytic, as these cells have relatively low mitochondrial content; however, more recently it has been demonstrated that certain neutrophil functions, including chemotaxis, NETosis and degranulation, require mitochondrial substrate oxidation (reviewed in ^35–37^).

The class I phosphoinositide 3-kinase (PI3K catalytic) subunits p110β/PIK3CB and p110γ/PIK3CG (Fig6A, B) were significantly less abundant in MS neutrophils compared to healthy controls. This major signalling pathway supports processes including phagocytosis ^38^ and NET formation ^39^. Interestingly, neutrophils from mice lacking PI3K subunits and associated proteins have reduced ROS and dysregulated degranulation ^40,41^, therefore reduced PI3K subunits in MS neutrophils could be linked to the reduction in ROS and altered degranulation observed.

**Figure 6:**
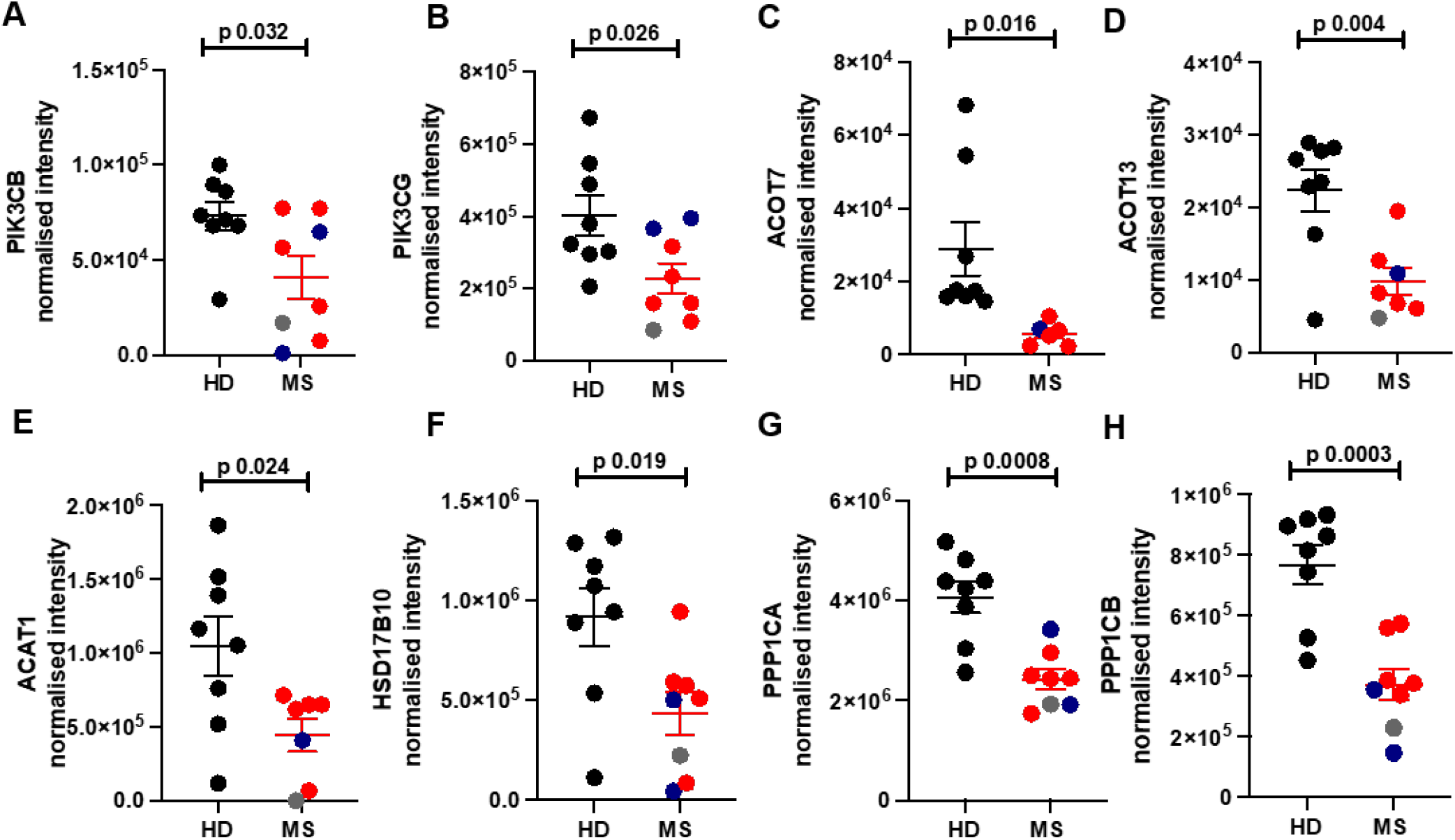
Reduced activity of metabolic pathways in MS neutrophils compared to healthy control cells. Peripheral blood was drawn from patients (RRMS, red, SPMS, blue or PPMS, grey) and healthy donors (HD, black). Neutrophils were isolated immediately via negative magnetic selection, centrifuged and cell pellets were snap-frozen. (A-F) Cell pellets were analysed via LC/MS for protein abundance. HD – healthy donor; MS – multiple sclerosis. N values: A-H –8. Statistical test: A-H – unpaired t test with Welch’s correction.

In addition, members of the acyl-CoA thioesterase (ACOT) family pathway, which regulate fatty acid metabolism, intracellular trafficking and storage of fatty acyl-CoAs ^42^, were strongly decreased in abundance (Fig6C, D).

Acetyl-CoA acetyltransferase 1 (ACAT1) and hydroxysteroid 17-beta dehydrogenase 10 (HSD17B10), involved in mitochondrial fatty acid beta oxidation, were also significantly downregulated (Fig6E, F).

Finally, two catalytic subunits of protein phosphatase 1 (PP1) (PPP1CA and PPP1CB), which regulates glycogen breakdown by dephosphorylation of rate limiting enzymes in this pathway ^43^ were also strongly reduced in MS neutrophils compared to healthy donors (Fig6G, H). This may be relevant, since glycogen stores were recently shown to critically support neutrophil survival and function ^43^. Taken together, these data indicate that MS neutrophils may demonstrate reduced activity of metabolic pathways including glycolysis, mitochondrial fatty acid oxidation (FAO) and glycogen breakdown. However, it will be important to define this at a functional level, particularly since certain of these proteins have complex roles in dynamic regulation of these pathways. For example, both impaired and excessive activity of the PI3K pathway, in context of loss- and gain-of-function subunit mutations, is associated with dysregulated metabolism and function of immune cells (reviewed in ^44^). The data may also reflect alterations in neutrophil maturation status, as immature neutrophils demonstrate higher mitochondrial content and dependency on lipid metabolism than mature counterparts (reviewed in ^45^).

### MS neutrophils are less able to suppress T cell activation

Neutrophils and T cells interact in the periphery, in lymph nodes, and in tissues including in the central nervous system ^12,13,46^. We and others have previously shown that interaction of resting neutrophils with T cells suppresses T cell proliferation and cytokine production ^8,47–49^, while contact with primed neutrophils, by contrast, promotes T cell activation ^47^. In particular, previous papers have indicated CD11b ^8^, cathelicidin ^11^, and elastase ^50^ can induce activation and differentiation of responding T cells, and these are all over-expressed in MS neutrophils in our dataset; in contrast, arginase is suppressive of T cells and this is not altered in our dataset. We therefore hypothesised that the increased activation of T cells seen in patients with MS may, in part, result from impaired tonic regulation by resting neutrophils, owing to the altered expression of T cell stimulating versus suppressive proteins.

We tested this hypothesis using a co-culture system we have previously established ^47^. Neutrophils and T cells were rapidly isolated from peripheral blood and cultured together for 24 hours at a 1:3 T cell: neutrophil ratio, in the presence of anti-CD3 antibodies. T cell phenotype was then assessed by flow cytometry. We compared MS neutrophil-MS T cell and HD neutrophil-HD T cell co-cultures and assessed activation of the T cells.

Here, co-culture with freshly isolated healthy neutrophils increased healthy T cell CD62L expression compared to T cells cultured alone (Fig7A-C), suggesting less activation of the T cells in response to the anti-CD3 stimulation. This was true for CD4^+^ and CD8^+^ T cells in these cultures. However, MS neutrophils did not significantly alter expression of CD62L on co-cultured T cells, indicating a reduced capacity to regulate T cell activity. Similarly, contact with healthy neutrophils also suppressed CD44 expression compared to T cells cultured alone (Fig7D,E), which, again, occurred to a much lower extent with MS neutrophils. Specifically, healthy neutrophils resulted in suppression of CD4^+^ T cell CD44 expression to 38.9% of original intensity, while MS neutrophils resulted in suppression only to 82.7% of original intensity.

**Figure 7:**
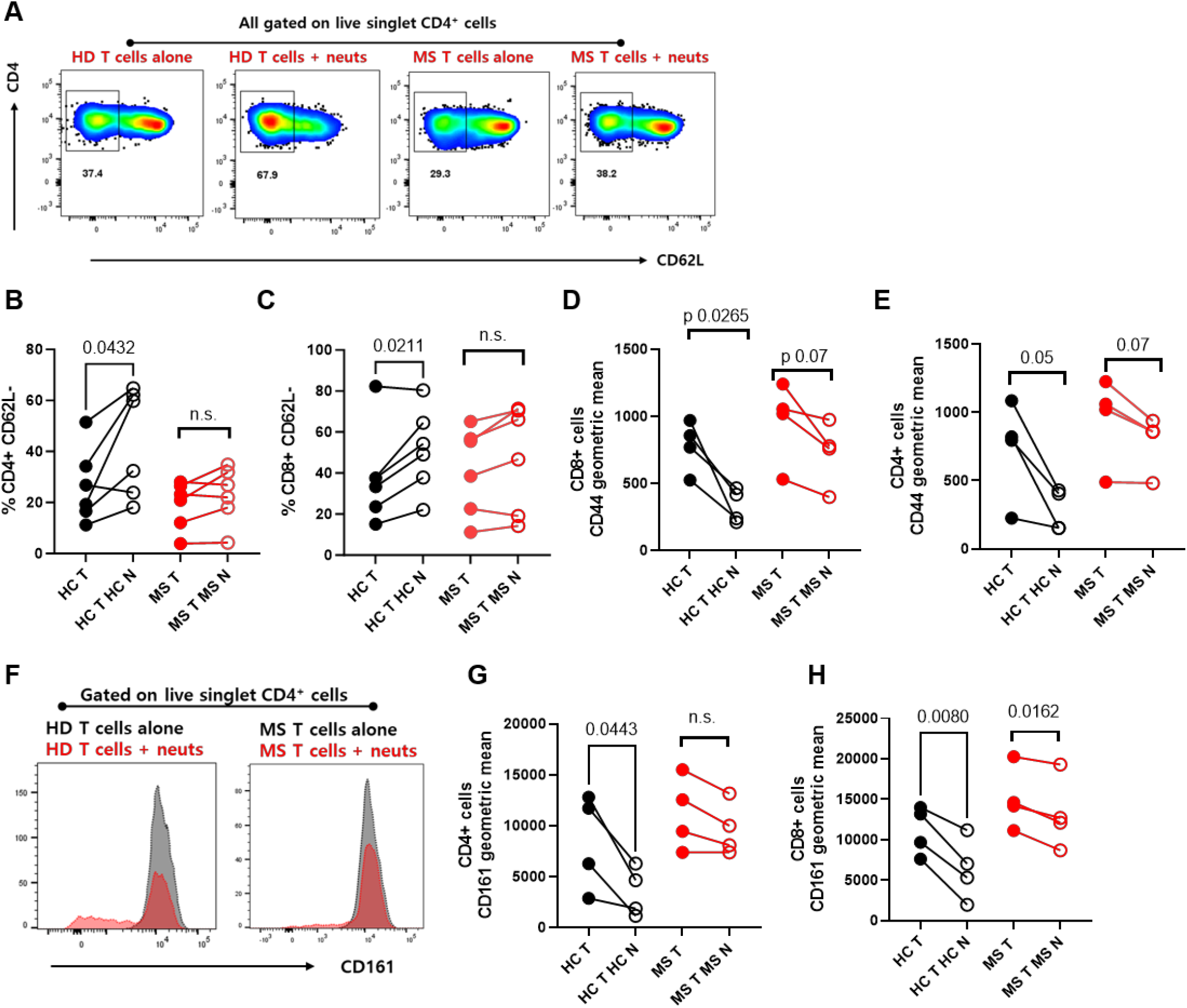
MS neutrophils are less able to suppress T cells than healthy donor cells. Peripheral blood was drawn from patients (MS) and healthy donors (HD). Neutrophils and total T cells were immediately isolated via negative magnetic selection. T cells were cultured either alone or with neutrophils (1:3 T:N) for 24 hours then assessed by flow cytometry. (A-E) activation of T cells was measured by CD62L and CD44 expression. (F-H) Expression of the Th17-related marker CD161 was quantified. N values: B, C – 6; D, E, G, H – 4. Statistical tests used – paired t test.

These data agree with previous studies which show that contact with resting neutrophils suppresses T cell activation, and extend these studies by demonstrating that this suppression is reduced with cells taken from MS patients.

### CD161 expression on T cells is altered following neutrophil contact

Of particular interest to the T cell phenotype in MS is expression of CD161. This surface molecule has shown to be a marker of IL-17-producing cells ^51^, and CD161^high^CD8^+^ cells are increased in the peripheral blood of MS patients ^52^, therefore we wondered whether neutrophils could regulate the expression of CD161 on T cells. After 24 hours′ incubation, CD4^+^ T cells incubated with HD neutrophils showed a significant suppression in CD161 expression on their surface compared to those activated but with no neutrophil contact (Fig7G). In contrast, those incubated with MS neutrophils were not significantly suppressed.

CD8^+^ T cells from both donors were suppressed by autologous neutrophils (Fig7H), although the MS neutrophils suppressed the CD161 expression less (to an average of 86% of their original expression) than the HD neutrophils did (to an average of 54% of their original expression). Therefore, it is possible that MS neutrophils are less able to suppress CD161 expression on T cells *in vivo*, leading to the presence of increased numbers of CD161^+^ cells.

## DISCUSSION

The aim of this study was to perform the first proteomic analysis of neutrophils from patients with Multiple Sclerosis, who are not on immune-modulating medication, to identify possible protein pathways that are dysregulated in MS neutrophils compared to healthy neutrophils.

We have shown here that neutrophils present in the circulation of MS patients are more activated and mature than in healthy controls, supporting previous studies performed using flow cytometry. One explanation for this could be that the increased inflammation and infectious burden in MS patients has triggered priming and activation of peripheral blood neutrophils. However, all of the patients showed increased maturity of the neutrophil populations, with very consistent magnitude of increase. This suggests it is very unlikely that viral infections are responsible for the increased maturity as it is unlikely all patients simultaneously were suffering from viral infections, of similar magnitude.

We have also shown that MS neutrophils have significantly increased granule protein content compared to HD neutrophils. Neutrophils degranulate in reverse order – that is, the secretory granules first, then tertiary granules are released, then the secondary and finally the primary granules. One explanation of our data is therefore that peripheral blood neutrophils in patients with MS contain higher levels of granule proteins, and also a decreased propensity to degranulate, so that only the tertiary granules have been released and the primary and secondary granules remain in the cells. However, the increased expression of CD63 on the surface of the MS neutrophils argues against this and suggest there is also increased release of primary granules by these cells. While we are unclear as to the mechanisms of release, this study clearly shows that multiple granule proteins are increased in MS neutrophils; as we know that granule proteins have a profound impact on the adaptive immune response ^53^, this is an important avenue to investigate. Previous studies in other conditions such as systemic lupus erythematosus have shown increased granule proteins in the serum of patients compared to healthy controls ^54,55^. However, this has been interpreted as increased degranulation by neutrophils, rather than the increased concentrations of these mediators to start with, which we see in our MS samples. We hypothesise that MS neutrophils have a disordered degranulation pattern compared to healthy controls as well as disordered production of granule proteins. A systematic analysis of combined neutrophil proteomics and secretomics in MS is now needed in a larger cohort of patients, to test this hypothesis and to understand the balance between degranulation and retention of granules.

A limitation of this study is that we isolated all neutrophils rapidly using negative magnetic selection rather than by density centrifugation. As a result, we could not separate cells according to their density, and low-and normal-density neutrophil populations will be combined in the proteomic analysis. Patients with MS have been observed to have a small but consistent population of low density neutrophils ^57^. The low-density population is small enough that it would be unlikely to explain large alterations in protein content across neutrophils as a whole, as we see here. However, it would be very interesting to repeat the granule protein analysis in particular in normal-density and low-density neutrophils, to understand whether there is one neutrophil population which is particularly dysregulated in disease.

We demonstrated decreased expression of some ROS proteins in MS neutrophils. Naegele and Hertwig have separately shown an increased functional respiratory burst in neutrophils from patients with MS ^17,22^. Of note, in both of these papers neutrophils were stimulated with fMLP and mean fluorescence intensity (MFl) of oxidized dihydrorhodamine was measured. Therefore, it is possible that increase in ROS is observed upon neutrophil stimulation only and not in neutrophils in a resting state (as the cells in this study were).

Our co-culture data demonstrates overall that healthy neutrophils suppress T cell activation, but this does not occur with cells from MS patients. It is possible that the altered composition of MS neutrophils results in different outcomes for T cells, or that the dysregulation of MS T cells means they cannot respond to neutrophils properly. The answer may, of course, be both of these. Neutrophils can express HLA-DR and present antigen to T cells ^58–62^, and so cross-donor cultures are unsuitable for unravelling the mechanisms behind the observations we make here. However, our data demonstrate that the neutrophil-T cell interaction is altered in MS compared to healthy donor cell cultures – a hugely interesting observation, as we are now aware of how important the interaction of neutrophils with T cells is for the longer term adaptive immune responses.

We also show that healthy neutrophils suppress the expression of CD161 on T cells in co-culture, but MS neutrophils were unable to do so. This supports previous data showing that patients with MS have higher numbers of these CD161^+^ cells ^52^. These cells are more likely to be IL-17 producing, suggesting that normal resting neutrophils have a role in suppressing differentiation of the Th17 subset of cells. In an added layer of complexity, this impact may be switched to a pro-inflammatory Th17-driving role when the neutrophils are primed, de-granulating or NETosing, as granule contents can induce Th17 differentiation^11,50^. We note also that T cell release of IL-17 – which occurs at higher levels in MS and is a key feature of the disease– is an inducer of granulopoiesis and of neutrophil maturation and activation ^63,64^. This may therefore be a positive feedback loop of increasing neutrophil maturation and IL-17 expression.

This study emphasises the role of neutrophils and their interactions with T cells in MS pathology, highlighting the importance of unravelling the complex interplay of these two cell types in longer term autoimmune disease.

## Acknowledgements

We are grateful to Fingerprints Proteomics, University of Dundee, to the Queen’s Medical Research Institute flow cytometry unit, University of Edinburgh (Shonna Johnston, Will Ramsey and Mari George) for assistance with experiments, and to the MS specialist nurses and patients in the clinic for the blood samples. We are also grateful to Dr Alejandro Brenes (University of Edinburgh) for critical reading of the manuscript.

## References

1. Harrington, L. E. et al. Interleukin 17–producing CD4+ effector T cells develop via a lineage distinct from the T helper type 1 and 2 lineages. Nat. Immunol. 6, 1123–1132 (2005).

2. Cua, D. J. et al. Interleukin-23 rather than interleukin-12 is the critical cytokine for autoimmune inflammation of the brain. Nature 421, 744–748 (2003).

3. Langrish, C. L. et al. IL-23 drives a pathogenic T cell population that induces autoimmune inflammation. J. Exp. Med. 201, 233–240 (2005).

4. Fletcher, J. M., Lalor, S. J., Sweeney, C. M., Tubridy, N. & Mills, K. H. G. T cells in multiple sclerosis and experimental autoimmune encephalomyelitis. Clin. Exp. Immunol. 162, 1–11 (2010).

5. Kunkl, M., Frascolla, S., Amormino, C., Volpe, E. & Tuosto, L. T Helper Cells: The Modulators of Inflammation in Multiple Sclerosis. Cells 9, 482 (2020).

6. Ando, D. G., Clayton, J., Kono, D., Urban, J. L. & Sercarz, E. E. Encephalitogenic T cells in the B10.PL model of experimental allergic encephalomyelitis (EAE) are of the Th-1 lymphokine subtype. Cell. Immunol. 124, 132–143 (1989).

7. Panitch, HillelS., Haley, AndreaS., Hirsch, RobertL. & Johnson, KennethP. EXACERBATIONS OF MULTIPLE SCLEROSIS IN PATIENTS TREATED WITH GAMMA INTERFERON. The Lancet 329, 893–895 (1987).

8. Pillay, J. et al. A subset of neutrophils in human systemic inflammation inhibits T cell responses through Mac-1. J. Clin. Invest. 122, 327–336 (2012).

9. Friedrich, M. et al. IL-1-driven stromal–neutrophil interactions define a subset of patients with inflammatory bowel disease that does not respond to therapies. Nat. Med. 27, 1970–1981 (2021).

10. Wright, H. L., Lyon, M., Chapman, E. A., Moots, R. J. & Edwards, S. W. Rheumatoid Arthritis Synovial Fluid Neutrophils Drive Inflammation Through Production of Chemokines, Reactive Oxygen Species, and Neutrophil Extracellular Traps. Front. Immunol. 11, (2021).

11. Minns, D. et al. The neutrophil antimicrobial peptide cathelicidin promotes Th17 differentiation. Nat. Commun. 12, 1285 (2021).

12. Smith, K. J. et al. The antimicrobial peptide cathelicidin drives development of experimental autoimmune encephalomyelitis in mice by affecting Th17 differentiation. PLoS Biol. 20, e3001554 (2022).

13. Lim, K. et al. Neutrophil trails guide influenza-specific CD8 + T cells in the airways. Science 349, aaa4352 (2015).

14. Patel, D. F. et al. Neutrophils restrain allergic airway inflammation by limiting ILC2 function and monocyte– dendritic cell antigen presentation. Sci. Immunol. 4, eaax7006 (2019).

15. Beauvillain, C. et al. Neutrophils efficiently cross-prime naive T cells in vivo. Blood 110, 2965–2973 (2007).

16. Vono, M. et al. Neutrophils acquire the capacity for antigen presentation to memory CD4+ T cells in vitro and ex vivo. Blood 129, 1991–2001 (2017).

17. Naegele, M. et al. Neutrophils in multiple sclerosis are characterized by a primed phenotype. J. Neuroimmunol. 242, 60–71 (2012).

18. Fahmi, R. M., Ramadan, B. M., Salah, H., Elsaid, A. F. & Shehta, N. Neutrophil-lymphocyte ratio as a marker for disability and activity in multiple sclerosis. Mult. Scler. Relat. Disord. 51, 102921 (2021).

19. Hemond, C. C., Glanz, B. I., Bakshi, R., Chitnis, T. & Healy, B. C. The neutrophil-to-lymphocyte and monocyte-to-lymphocyte ratios are independently associated with neurological disability and brain atrophy in multiple sclerosis. BMC Neurol. 19, 23 (2019).

20. Demirci, S., Demirci, S., Kutluhan, S., Koyuncuoglu, H. R. & Yurekli, V. A. The clinical significance of the neutrophil-to-lymphocyte ratio in multiple sclerosis. Int. J. Neurosci. 1–7 (2015) doi:10.3109/00207454.2015.1050492.

21. Aubé, B. et al. Neutrophils Mediate Blood–Spinal Cord Barrier Disruption in Demyelinating Neuroinflammatory Diseases. J. Immunol. 193, 2438–2454 (2014).

22. Hertwig, L. et al. Distinct functionality of neutrophils in multiple sclerosis and neuromyelitis optica. Mult. Scler. J. 22, 160–173 (2016).

23. Rissanen, E., Remes, K. & Airas, L. Severe neutropenia after rituximab-treatment of multiple sclerosis. Mult. Scler. Relat. Disord. 20, 3–5 (2018).

24. Burt, R. et al. Collection of hematopoietic stem cells from patients with autoimmune diseases. Bone Marrow Transplant. 28, 1–12 (2001).

25. Steinbach, K., Piedavent, M., Bauer, S., Neumann, J. T. & Friese, M. A. Neutrophils Amplify Autoimmune Central Nervous System Infiltrates by Maturing Local APCs. J. Immunol. 191, 4531–4539 (2013).

26. Rumble, J. M. et al. Neutrophil-related factors as biomarkers in EAE and MS. J. Exp. Med. 212, 23–35 (2015).

27. Long, M. B. et al. Extensive acute and sustained changes to neutrophil proteomes post-SARS-CoV-2 infection. Eur. Respir. J. 2300787 (2023) doi:10.1183/13993003.00787-2023.

28. Hoogendijk, A. J. et al. Dynamic Transcriptome-Proteome Correlation Networks Reveal Human Myeloid Differentiation and Neutrophil-Specific Programming. Cell Rep. 29, 2505–2519.e4 (2019).

29. Reyes, L. et al. A type I IFN, prothrombotic hyperinflammatory neutrophil signature is distinct for COVID-19 ARDS--. Wellcome Open Res. 6, 38 (2021).

30. Ghaderi, S. et al. Hospitalization following influenza infection and pandemic vaccination in multiple sclerosis patients: a nationwide population-based registry study from Norway. Eur. J. Epidemiol. 35, 355–362 (2020).

31. Lechner-Scott, J., Waubant, E., Levy, M., Hawkes, C. & Giovannoni, G. Is multiple sclerosis a risk factor for infections? Mult. Scler. Relat. Disord. 41, 102184 (2020).

32. Vinogradova, Y., Hippisley-Cox, J. & Coupland, C. Identification of new risk factors for pneumonia: population-based case-control study. Br. J. Gen. Pract. 59, e329–e338 (2009).

33. Kuijpers, T. W. et al. Membrane surface antigen expression on neutrophils: a reappraisal of the use of surface markers for neutrophil activation. Blood 78, 1105–1111 (1991).

34. Ohayon, D. et al. Cytosolic PCNA interacts with p47phox and controls NADPH oxidase NOX2 activation in neutrophils. J. Exp. Med. 216, 2669–2687 (2019).

35. Injarabian, L., Devin, A., Ransac, S. & Marteyn, B. S. Neutrophil Metabolic Shift during Their Lifecycle: Impact on Their Survival and Activation. Int. J. Mol. Sci. 21, 287 (2019).

36. Cao, Z. et al. Roles of mitochondria in neutrophils. Front. Immunol. 13, 934444 (2022).

37. Kumar, S. & Dikshit, M. Metabolic Insight of Neutrophils in Health and Disease. Front. Immunol. 10, 2099 (2019).

38. Pan, T. et al. Immune effects of PI3K/Akt/HIF-1α-regulated glycolysis in polymorphonuclear neutrophils during sepsis. Crit. Care 26, 29 (2022).

39. Rodríguez-Espinosa, O., Rojas-Espinosa, O., Moreno-Altamirano, M. M. B., López-Villegas, E. O. & Sánchez-García, F. J. Metabolic requirements for neutrophil extracellular traps formation. Immunology 145, 213–224 (2015).

40. Gambardella, L. et al. The GTPase-activating protein ARAP3 regulates chemotaxis and adhesion-dependent processes in neutrophils. Blood 118, 1087–1098 (2011).

41. Kulkarni, S. et al. PI3Kβ plays a critical role in neutrophil activation by immune complexes. Sci. Signal. 4, ra23 (2011).

42. Tillander, V., Alexson, S. E. H. & Cohen, D. E. Deactivating Fatty Acids: Acyl-CoA Thioesterase-Mediated Control of Lipid Metabolism. Trends Endocrinol. Metab. TEM 28, 473–484 (2017).

43. Sadiku, P. et al. Neutrophils Fuel Effective Immune Responses through Gluconeogenesis and Glycogenesis. Cell Metab. 33, 411–423.e4 (2021).

44. Lucas, C. L., Chandra, A., Nejentsev, S., Condliffe, A. M. & Okkenhaug, K. PI3Kδ and primary immunodeficiencies. Nat. Rev. Immunol. 16, 702–714 (2016).

45. Jeon, J.-H., Hong, C.-W., Kim, E. Y. & Lee, J. M. Current Understanding on the Metabolism of Neutrophils. Immune Netw. 20, e46 (2020).

46. Beauvillain, C. et al. CCR7 is involved in the migration of neutrophils to lymph nodes. Blood 117, 1196–1204 (2011).

47. Minns, D., Smith, K. J., Hardisty, G., Rossi, A. G. & Gwyer Findlay, E. The Outcome of Neutrophil-T Cell Contact Differs Depending on Activation Status of Both Cell Types. Front. Immunol. 12, 633486 (2021).

48. Mensurado, S. et al. Tumor-associated neutrophils suppress pro-tumoral IL-17+ γδ T cells through induction of oxidative stress. PLOS Biol. 16, e2004990 (2018).

49. Germann, M. et al. Neutrophils suppress tumor-infiltrating T cells in colon cancer via matrix metalloproteinase-mediated activation of TGF β. EMBO Mol. Med. 12, e10681 (2020).

50. Souwer, Y. et al. Human TH17 cell development requires processing of dendritic cell–derived CXCL8 by neutrophil elastase. J. Allergy Clin. Immunol. 141, 2286–2289.e5 (2018).

51. Maggi, L. et al. CD161 is a marker of all human IL-17-producing T-cell subsets and is induced by RORC. Eur. J. Immunol. 40, 2174–2181 (2010).

52. Annibali, V. et al. CD161highCD8+T cells bear pathogenetic potential in multiple sclerosis. Brain 134, 542–554 (2011).

53. Minns, D., Smith, K. J. & Findlay, E. G. Orchestration of Adaptive T Cell Responses by Neutrophil Granule Contents. Mediators Inflamm. 2019, 8968943 (2019).

54. Sthoeger, Z. M., Bezalel, S., Chapnik, N., Asher, I. & Froy, O. High α-defensin levels in patients with systemic lupus erythematosus. Immunology 127, 116–122 (2009).

55. Vordenbäumen, S. et al. Elevated levels of human beta-defensin 2 and human neutrophil peptides in systemic lupus erythematosus. Lupus 19, 1648–1653 (2010).

56. Sengeløv, H. et al. Mobilization of granules and secretory vesicles during in vivo exudation of human neutrophils. J. Immunol. 154, 4157–4165 (1995).

57. Ostendorf, L. et al. Low-Density Granulocytes Are a Novel Immunopathological Feature in Both Multiple Sclerosis and Neuromyelitis Optica Spectrum Disorder. Front. Immunol. 10, 2725 (2019).

58. Iking-Konert, C. et al. Polymorphonuclear neutrophils in Wegener’s granulomatosis acquire characteristics of antigen presenting cells. Kidney Int. 60, 2247–2262 (2001).

59. Pliyev, B. K., Dimitrieva, T. V. & Savchenko, V. G. Cytokine-mediated induction of MHC class II in human neutrophils is dependent on NADPH oxidase activity. Eur. J. Cell Biol. 94, 67–70 (2015).

60. Reinisch, W. et al. Donor dependent, interferon-γ induced HLA-DR expression on human neutrophils in vivo. Clin. Exp. Immunol. 133, 476–484 (2003).

61. Fanger, N. A. et al. Activation of Human T Cells by Major Histocompatability Complex Class II Expressing Neutrophils: Proliferation in the Presence of Superantigen, But Not Tetanus Toxoid. Blood 89, 4128–4135 (1997).

62. Moffat, A. & Gwyer Findlay, E. Evidence for antigen presentation by human neutrophils. Blood J. blood.2023023444 (2024) doi:10.1182/blood.2023023444.

63. Chen, X. et al. Reactive granulopoiesis depends on T-cell production of IL-17A and neutropenia-associated alteration of gut microbiota. Proc. Natl. Acad. Sci. 119, e2211230119 (2022).

64. Deshmukh, H. S. et al. The microbiota regulates neutrophil homeostasis and host resistance to Escherichia coli K1 sepsis in neonatal mice. Nat. Med. 20, 524–530 (2014).

